# rdmc: an open source R package implementing convergent adaptation models of Lee and Coop (2017)

**DOI:** 10.1101/2020.04.22.056150

**Authors:** Silas Tittes

**Author notes:** Dept. of Evolution and Ecology, University of California, Davis, CA, USA.

## Abstract

The availability of whole genome sequencing data from multiple related populations creates opportunities to test sophisticated population genetic models of convergent adaptation. Recent work by Lee and Coop (2017) developed models to infer modes of convergent adaption at local genomic scales, providing a rich framework for assessing how selection has acted across multiple populations at the tested locus. Here I present, *rdmc*, an R package that builds on the existing software implementation of Lee and Coop (2017) that prioritizes ease of use, portability, and scalability. I demonstrate installation and comprehensive overview of the package’s current utilities.

## Introduction

Convergent adaptation occurs when natural selection independently orchestrates the evolution of the same set of trait or traits in multiple populations (Takuno *et al.* 2015; Tishkoff *et al.* 2007; Yeaman *et al.* 2016; Losos 2011). Efforts by Lee and Coop (2017) used coalescent theory to develop composite likelihood models to infer which among several competing modes of convergent adaptation best explains allele frequencies at a putatively selected region. These models provide rich statistical information about the inferred adaptive mutation, including its location along the region, the strength of selection, migration rate, age, and its initial allele frequency prior to selection.

To facilitate use of the convergent adaptation models of Lee and Coop (2017), I developed rdmc, an R package implementing their models that was designed to be easy to use, portable, and scalable. In this short manuscript, I provide an overview of the usage and installation of the package, concluding with opportunities for future improvements and expansion to the software.

## Materials and Methods

Lee and Coop (2017) described three distinct modes of convergent adaption: independent mutations, where two or more populations independently gain the selected mutation; migration, where the mutation occurs once and subsequently migrates to multiple populations prior to fixation; and standing variation, where the mutation was present at low frequency in the ancestral population prior to divergence. The models are composite likelihood-based, where likelihood calculations are made over a grid of user-chosen input parameters.

### Data requirements

Using *rdmc* requires two kinds of allele frequency data. The first is allele frequencies from unlinked neutral sites across all populations. The second is allele frequencies from at least three populations that have putatively undergone convergent adaptation at a specific locus, and three or more populations that did not. Sample allele frequencies can be estimated with a number of existing software resources including VCFtools (Danecek et al. 2011) and ANGSD (Korneliussen et al. 2014). Typically, the allele frequencies at sites that have putatively undergone convergent adaptation will have been identified prior to using *rdmc.* Numerous methods exists for identifying such regions, such as finding overlapping selective sweeps in multiple populations (Stetter *et al.* 2020), or by identifying regions with elevated *F_ST_* values between populations that putatively did and did not experience convergent adaptation (Hohenlohe *et al.* 2010). Additionally, *rdmc* requires an estimate of the per base recombination rate for the region or regions of interest, and an estimate of the effective population size. Assuming a mutation rate, effective population size can be estimated from genetic diversity (Gillespie 2004), or inferred via multiple methods (Gutenkunst *et al.* 2010; Schiffels and Durbin 2014; Excoffier and Foll 2011). Likewise, local recombination rates can be derived from genetic maps (Swarts *et al.* 2014), or inferred (Chan *et al.* 2012; McVean and Auton 2007; Adrion *et al.* 2020). At the time of writing, all available models in *rdmc* assume a single effective population size for all populations. Depending on which modes of convergent adaptation are being investigated, users must also provide vectors of selection coefficients, migration rates, allele frequency ages, and initial allele frequencies prior to selection. The exhaustive list of required inputs and their definitions is given in Table 1.

**Table 1.** Description of the arguments used with the function *parameter_barge()*.

| Function argument | Description |
| --- | --- |
| neutral_freqs | Matrix of allele frequencies at putatively neutral sites with dimensions, number of populations x number of sites. |
| selected_freqs | Matrix of allele frequencies at putatively selected sites with dimensions, number of populations x number of sites. |
| selected_pops | Vector of indices for populations that experienced selection. |
| positions | Vector of genomic positions for the selected region. |
| n_sites | Integer for the number of sites to propose as the selected site. Sites are uniformly placed along positions using `seq(min(positions), max(positions), length.out = n_sites)`. Must be less than or equal to `length(positions)`. Cannot be used with `sel_sites`. |
| sel_sites | Optional vector of sites to propose as selected site. Useful if particular sites are suspected to be under selection. Cannot be used with `n_sites`. |
| sample_sizes | Vector of sample sizes of length number of populations. (i.e. twice the number of diploid individuals sampled in each population). |
| num_bins | The number of bins in which to bin alleles a given distance from the proposed selected sites. |
| sets | A list of population indices, where each element in the list contains a vector of populations with a given mode of convergence. For example, if populations 2 and 6 share a mode and population 3 has another, `sets = list(c(2,6), 3)`. Only required for fitting models with mixed modes. Must be used in conjunction with the "modes" argument. |
| modes | Character vector of length sets defining mode for each set of selected populations ("independent", "standing", and/or "migration"). Only required for fitting models with mixed modes. More details about the modes is available on help page for `mode_cle`. |
| sels | Vector of proposed selection coefficients. |
| migs | Vector of proposed migration rates (proportion of individuals of migrant origin each generation). Cannot be 0. |
| times | Vector of proposed times in generations the variant is standing in populations before selection occurs and prior to migration from source population. |
| gs | Vector of proposed frequencies of the standing variant. |
| sources | Vector of population indices to propose as the source population of the beneficial allele. Used for both the migration and standing variant with source models. Note: the source must be one of the populations contained in `selPops`. |
| Ne | Effective population size (assumed equal for all populations). |
| rec | Per base recombination rate for the putatively selected region. |
| locus_name | String to name the locus. Helpful if multiple loci will be combined in subsequent analyses. Defaults to "locus". |
| cholesky | Logical to use cholesky factorization of covariance matrix. Used for both inverse and determinant. Faster, but not guaranteed to work for all data sets. TRUE by default. if FALSE, `ginv` from MASS is used. |

### Installation and dependencies

Installation of *rdmc* requires the r package *devtools* (Wickham *et al.* 2020b). With devtools available, the package can be installed and made locally available with the following R commands:

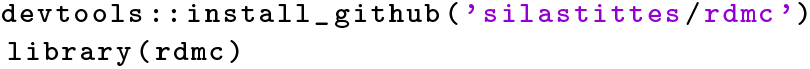

In addition to *devtools, rdmc* depends on several other packages. Namely, *MASS* (Venables and Ripley 2002), *dplyr* (Wickham *et al.* 2020a), *tidyr* (Wickham and Henry 2020), *purrr* (Henry and Wickham 2019), *magrittr* (Bache and Wickham 2014), and *rlang* (Henry and Wickham 2020). All dependencies are automatically installed or updated when the installation command above is issued. I encourage users to update to the most recent version of R prior to issuing any of the above commands.

### Specifying parameters and input data

For convenience, the original simulated example data generated by Lee and Coop (2017) are provided with the installation and can be loaded with:

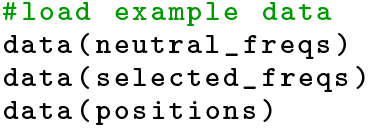

The example data consists of 10,000 simulated base pairs from six populations, three of which (with indices 1,3,5) independently mutated to the selected allele at position 0, along with three other populations that evolved neutrally. Allele frequencies must be be passed to *rdmc* as a matrix, where each row is a population and each column is a locus. Users should note that the simulated positions here take on values between zero and one, but that base pair positions along the chromosomes of empirical data should not be altered prior to fitting the models.

When fitting possible convergent adaptation models, several quantities are reused regardless of which modes of convergent adaptation are of interest. In efforts to minimize computation, all parameters and quantities that are common across models are stored in a single named list generated with the function *parameter_barge()* that can be used when fitting any of the possible models. The list of quantities is generated using:

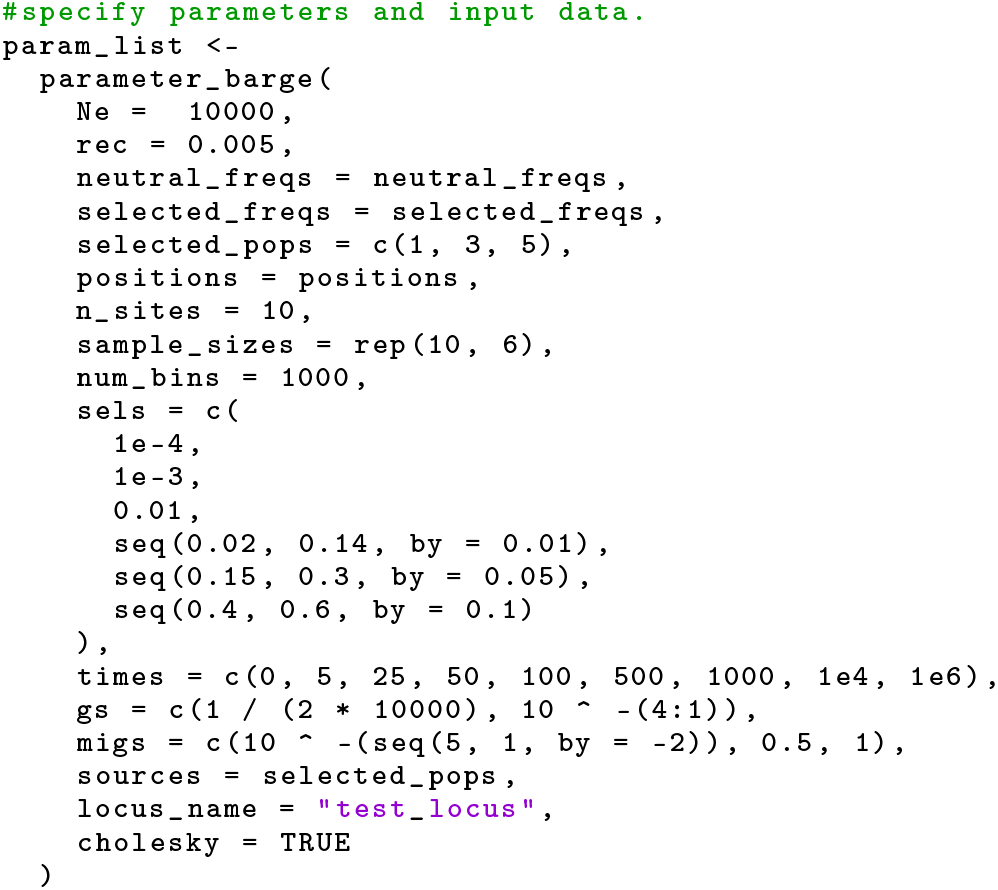

where all the arguments are fully described in Table 1. This command also determines the grid of parameter values (namely the arguments, *sels, times, gs, migs, sources*, and *n_sites* or *positions*) that will be used in the likelihood calculations. Depending on which modes of convergent adaptation are being studied, some of these arguments may not be used for inferences. Users must still input values for all arguments. Missing values for parameters (*NA*) can be used, as long as that parameter is not used.

Naturally, features of the input data (the density and amount of variation in the allele frequencies, the effective population size, and the mutation and recombination rates), will impact the model results, and will determine the resolution we have to infer the model parameters. The number and density of points along the grid of parameters also effect the resolution one has to make inferences. However, computation time and memory usage may become infeasible if these grids are made too large.

### Fitting the models

Once the parameter barge is constructed, all models can be fit using this list of quantities as the only data input. All of the mode types (neutral, independent mutations, standing variation with and without a source population, migration, and mixed-modes) are implemented using the same function, *mode_cle()*, passing the desired mode as an argument to the function. The neutral, independent mutations, migration, and standing variation with a source population modes can be fit, respectively with:

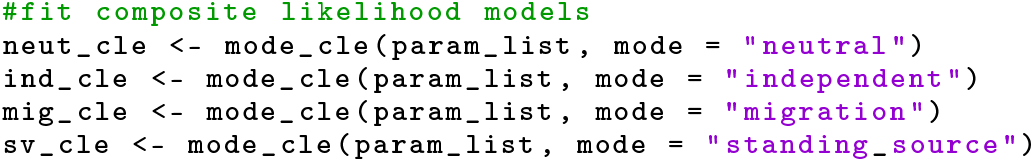

Models of mixed modes, where specified populations are modeled under different modes, can be also implemented by modifying the parameter list object in-place. Specifically, mixed modes are constructed by adding the *sets* and *modes* arguments, which groups the population indices according the vector of modes, and specifies which modes are to be used. For example, to fit a model where populations with indices 1 and 3 adapted via standing variation, and population 5 gained the same mutation independently, and another mixed-mode model where populations 1 and 3 adapted via migration, and population 5 mutated independently:

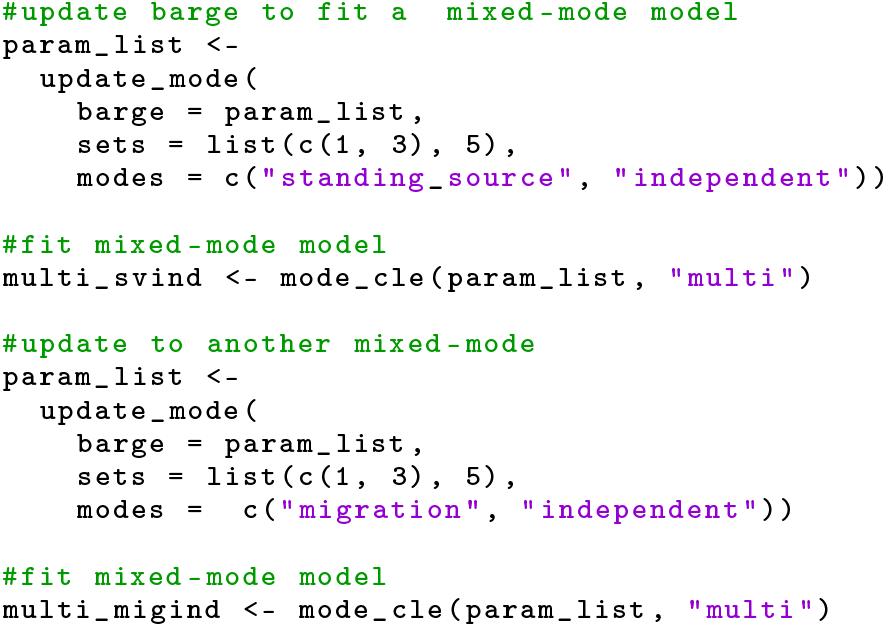

Regardless of which mode is used when calling *mode_cle()*, the data frame returned will always contain the same 10 features: The 6 grid parameters generated by *parameter_barge()* (Table 1), the composite likelihood score calculated over all possible combinations of the grid parameters, the indices of the selected populations, and the names of the locus and mode that were used. To always maintain the same number of columns, missing (*NA*) values are added when variables are not used for a given mode type. As will be shown below, this design facilitates combining results from multiple models for downstream analyses.

## Results and Discussion

### Benchmarking

The computation time and memory usage of *rdmc* increases with the complexity of the model and size of the input data used. Compared to the original code implemented by Lee and Coop (2017), *rdmc* is slightly slower computationally, but requires substantially less memory. However, the reduced memory allocation for *rdmc* only occurs when Cholesky factorization is used to obtain the inverse of the neutral and selected covariance matrices (Tables 1 and 2). Alternatively, the matrix inverses are obtained using *ginv()* from the MASS package (Venables and Ripley 2002), which requires a larger memory allocation, but will still approximate the inverse even if the covariance matrix is not positive definite. Users are therefore encouraged to use the default *parameter_barge()* argument *cholesky = TRUE.* If the Cholesky factorization fails, an informative error message will be generated encouraging setting *cholesky = FALSE.*

The composite likelihood calculations are made over a grid of input parameters chosen when constructing the parameter barge (code shown above), hence, a denser grid will also have a considerable impact on time and memory usage. The size of the example data provided gives a realistic sense of memory and time usage for potential empirical data. While most modern laptops are capable of handling the required memory, many users will be interested in genome-wide analysis, where the mode of convergence for many separate regions are of interest. In these instances, cloud or high performance computing environments will be more appropriate. Making *rdmc* a portable and easy to install R package simplifies running separate genomic regions as independent jobs in parallel using workflows such as Snakemake (Köster and Rahmann 2012) or Nextflow (Di Tommaso *et al.* 2017).

**Table 2.**
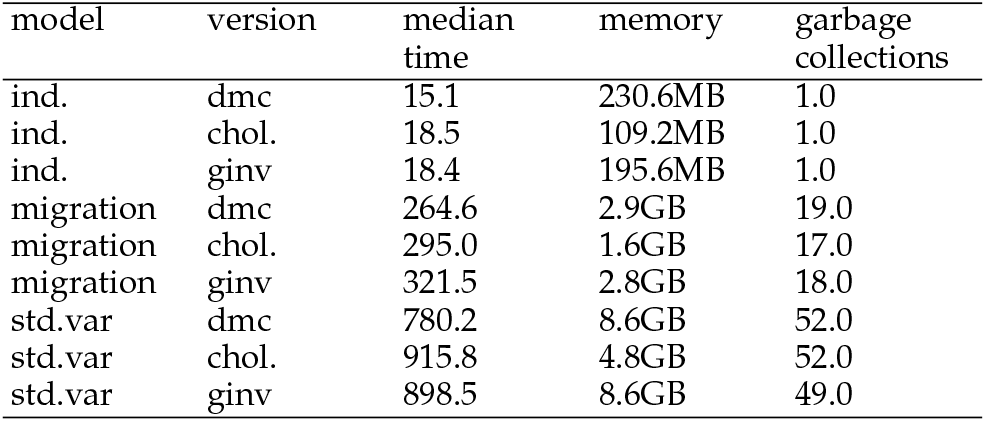
Benchmarking of three *rdmc* model types. Computation time, memory allocation, and the number of garbage collections are reported for the original (dmc) code written by Lee and Coop (2017), and the two matrix inversion methods available in *rdmc* (ginv and chol.). Median time was estimated using 5 iterations of each model. Execution time is reported in seconds. Benchmarking was conducted using the R package, *bench* (Hester 2020). Code was executed in an interactive job on the UC Davis Farm HPC (2.00GHz Intel Xeon CPU, 124GB RAM).

| model | version | median time | memory | garbage collections |
| --- | --- | --- | --- | --- |
| ind. | dmc | 15.1 | 230.6MB | 1.0 |
| ind. | chol. | 18.5 | 109.2MB | 1.0 |
| ind. | ginv | 18.4 | 195.6MB | 1.0 |
| migration | dmc | 264.6 | 2.9GB | 19.0 |
| migration | chol. | 295.0 | 1.6GB | 17.0 |
| migration | ginv | 321.5 | 2.8GB | 18.0 |
| std.var | dmc | 780.2 | 8.6GB | 52.0 |
| std.var | chol. | 915.8 | 4.8GB | 52.0 |
| std.var | ginv | 898.5 | 8.6GB | 49.0 |

### Extracting useful quantities and visualization

Once the models of interest have finished, the common format of the returned data frames allows all of the inferences to be combined into a single data frame, which simplifies creation of statistical and graphical summaries, and storage:

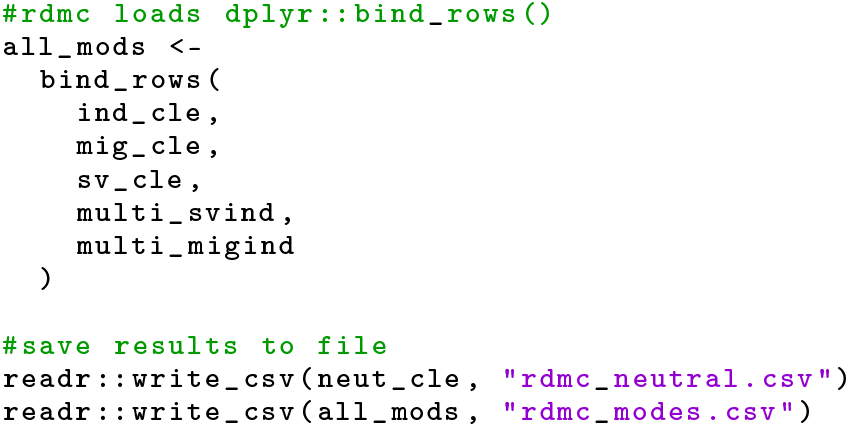

With a single data frame containing output from all tested models, there are many visualization and summary methods are available in the R ecosystem (R Core Team 2020). For example, the maximum composite likelihood estimate of the selection coefficient for each model can be accessed with:

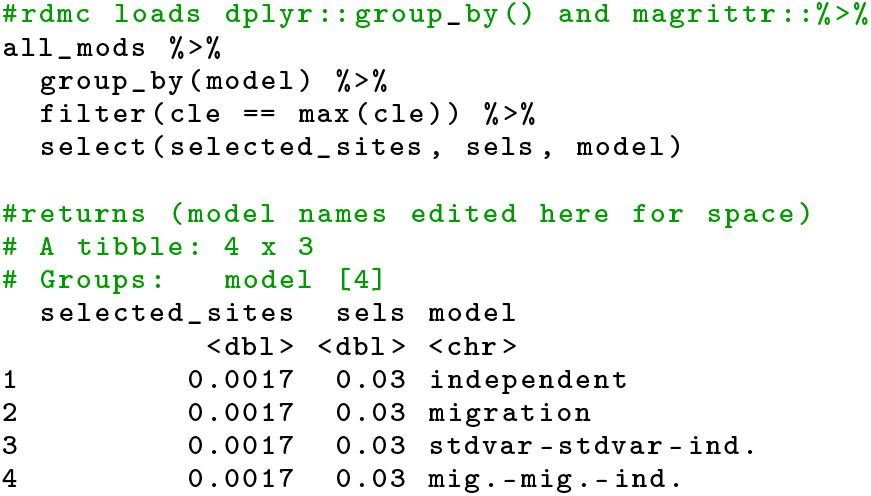

Visualizing the composite likelihood values by genomic position (relative to the neutral composite likelihood) (Figure 1) can be done with:

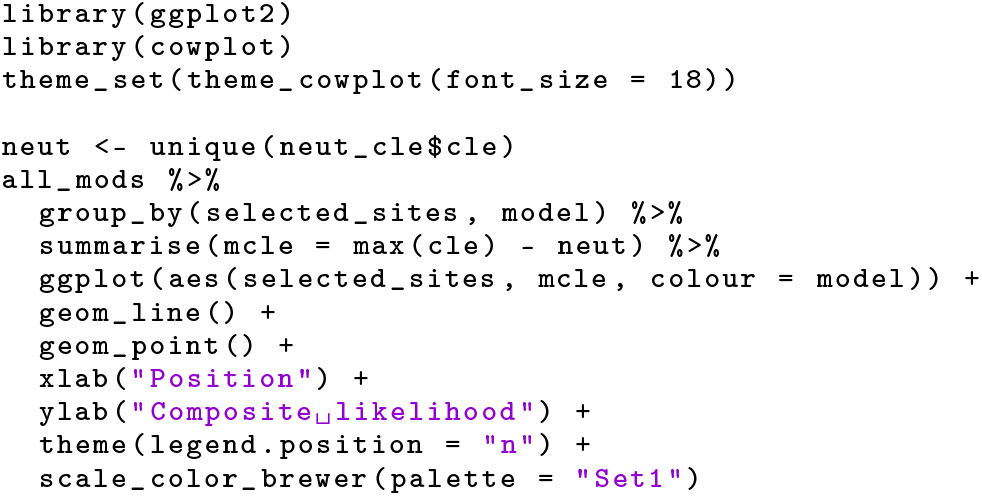

**Figure 1.**
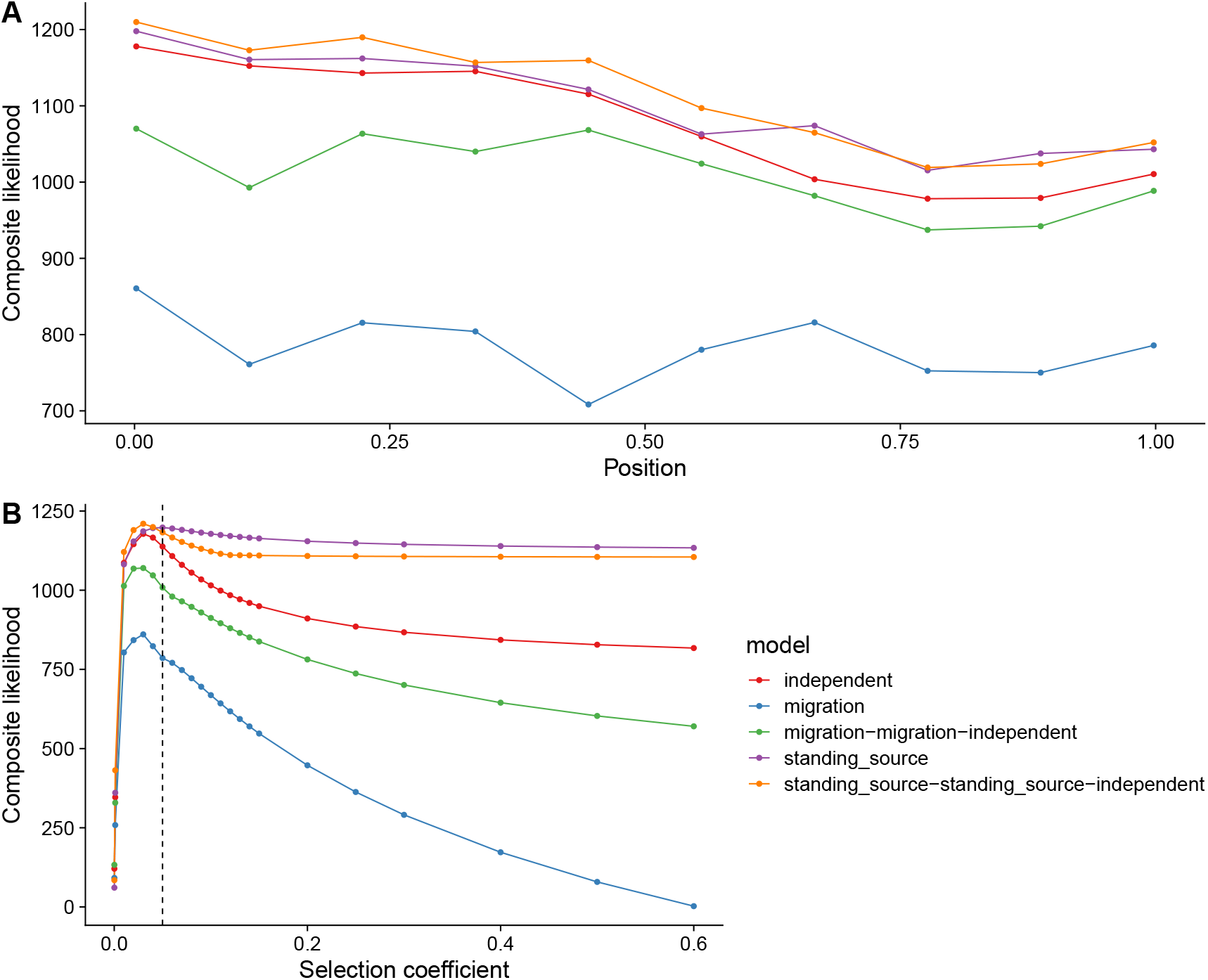
Visualizing rdmc results for several modes. (A) The composite likelihood score at each of the 10 proposed sites of selection for each model. The true selected site was modeled at position 0. The data was simulated as independent mutations in the three selected populations. (B) The composite likelihood scores over grid of selection coefficients. Dotted line indicates true selection coefficient (s = 0.05) the data was modeled under. Visualizations were made using the R packages, *ggplot2* (Wickham 2016) and *cowplot* (Wilke 2019)

Lastly, one can visualize the likelihood surface with respect to specific parameter, such as selection (Figure 1):

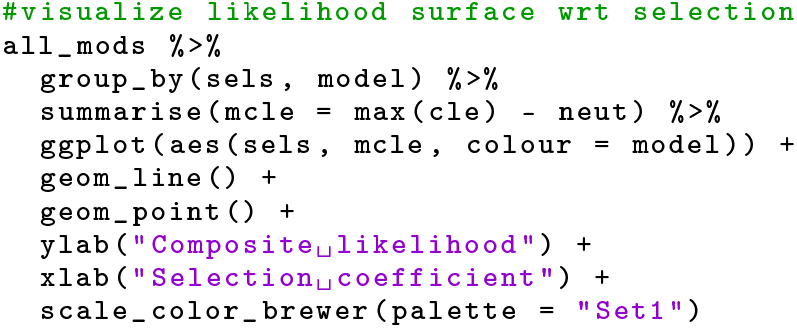

### Concluding remarks and future developments

*rdmc* was made to facilitate the use of convergent adaptation models of Lee and Coop (2017). The package is easy to install, and requires only a few lines of code to generate and analyze the output. By making *rdmc* an R package, the code is highly portable and has relatively few, highly maintained dependencies, making it simpler to adopt to different computing systems. Because of its portability and ease of use, *rmdc* also simplifies downstream tasks which facilitates usage at large scales, such as modeling thousand of genomic regions simultaneously on high performance computing resources.

Several elaborations to the currently available utilities in the *rdmc* package could be addded. Since the methods developed in Lee and Coop (2017), additional models have been developed, including ones that can use putatively selected deletion variation, strong selection, concurrent sweeps, and variation in population size among populations (Oziolor *et al.* 2019). Lee and Coop (2017) also introduced parametric bootstrapping to evaluate support for alternative modes. While not currently incorporated into *rdmc*, future development of the package would include functions to perform bootstrapping. However, for the same reasons mentioned above, *rdmc* should facilitate creation and computation of bootstrap replicates in parallel.

## Web resources

The source of the package and workflow outlined above are available at https://github.com/silastittes/rdmc. The package is released under GNU General Public License (v3.0). All of the presented analyses were computed on a personal laptop (x86_64, Apple) using R version 4.0.0 2020-04-24).

The original code associated with Lee and Coop (2017) is available https://github.com/kristinmlee/dmc.

## Acknowledgments

I would like to thank Jeff Ross-Ibarra for early reviews and encouragement to write this manuscript, to Kristin Lee, Sivan Yair, and Graham Coop for developing the methods and code that form the foundation of the *rdmc* package, and for giving me their blessing in pursuing the development of the package. Lastly, I thank Felix Andrews for helpfully suggesting the function name, *parameter_barge().*

